# Developing terrestrial environmental DNA sampling methods for detecting arboreal invasive reptiles: a case study of the green anole in the Ogasawara Island, Japan

**DOI:** 10.1101/2025.05.24.655731

**Authors:** Satsuki Tsuji, Yuki Murakami, Mitsuhiko Toda, Yosuke Yagami, Kou Ashizawa, Takuro Nishiwaki, Nayuta Yamamoto

## Abstract

Early detection of invasive alien species is essential for preventing establishment and mitigating ecological impacts, particularly in island ecosystems harbouring evolutionarily isolated endemic species. Recently, despite increasing reptile introductions and their suggested widespread impacts, methods for monitoring arboreal invasive reptiles remain limited. This study addressed the need for practical detection tools by developing terrestrial environmental DNA (eDNA) sampling methods that collect DNA from leaf surfaces to detect the arboreal invasive green anole (*Anolis carolinensis*) in the Ogasawara Islands, Japan. Two methods, wiping leaf surfaces with gauze and rinsing with sprayed water, were tested. At an invaded site, green anoles were successfully detected in eight out of 10 samples in both methods. Although no significant difference in eDNA concentration was observed, the wiping method was selected for its greater simplicity. Subsequently, the relationship between green anole population density and eDNA concentrations detected using the wiping method was investigated, suggesting a significant positive relationship. This is the first report demonstrating that terrestrial eDNA concentrations can reflect arboreal reptile population density, suggesting potential applications in quantitative terrestrial biodiversity assessments. Furthermore, the successful detection of eDNA even in the extremely low-density habitat of the green anole demonstrates the usefulness of eDNA-based surveys for early detection of invasions. The method developed here may be broadly applicable to terrestrial biodiversity monitoring, especially in tree-dwelling taxa. Given accelerating biological invasions and biodiversity loss, this approach is expected to benefit managers, conservationists, and researchers concerned with terrestrial ecosystems.

## 1 Introduction

In recent years, there has been an increasing need to monitor invasive alien species closely. Biological invasions are usually virtually irreversible and are one of the main causes of severe negative impacts on ecosystems (Clavero and García-Berthou, 2005; Diagne et al., 2021; Mack et al., 2000; Vitousek et al., 1997). Particularly on oceanic islands that have never been connected to a large landmass since their formation and are thus endemic species hotspots, evolutionarily isolated native species are often vulnerable to introduced predators and competitors. Invasive species have been implicated in 86% of recorded extinctions of island endemics since 1500 A.D. (Bellard et al., 2016; Spatz et al., 2017). Although invasive species originate from a wide range of taxonomic groups, there are substantial biases in the taxa that have been the focus of detailed case studies (e.g. top four: plants 44%, insects 18%, fish 8%, crustaceans 8%; Pyšek et al., 2008). Reptiles, one of the least studied taxa, account for only 2% of invasion research when combined with amphibians (Pyšek et al., 2008). However, invasive reptiles have been shown to exert ecological impacts through various mechanisms (Kraus, 2015). Nevertheless, records of the initial introductions of reptiles have increased exponentially since at least the 1950s, likely due to expanding transport networks and the growth of online trade (Seebens et al., 2017).

There is now an urgent need to develop simple and efficient methods for detecting invasive alien species of reptiles in the early stages of their invasion. Biological invasions require management strategies that incorporate both prevention and early response, with the success of early detection monitoring (EDM) being particularly crucial to prevent establishment and minimise impacts (Kaiser and Burnett, 2010; G. S. Peterson et al., 2022). EDM requires detection at the early invasion stage when target species are still few and localised, making it inherently challenging and requiring a balance between limited resources and comprehensive sampling efforts (Trebitz et al., 2017). However, EDM for reptiles is particularly challenging, as they typically hide in trees, behind rocks or under decaying leaves, with their ectothermic physiology further complicating detection. One of the invasive lizards, green anole (*Anolis carolinensis*), exemplifies such challenges. This species is an arboreal and insectivorous lizard native to North America. They have invaded several island environments, including the Hawaiian Islands, the Mariana Islands, the Ogasawara Islands and the Okinawa Islands in Japan, seriously affecting native ecosystems (Karube, 2010). Particularly in the Ogasawara Islands, it is said that their distribution range has expanded between islands since its first invasion in the 1960s (Sugawara et al., 2015; Toda et al., 2010). On islands in the early stages of invasion or those not yet invaded, EDM efforts led by the Ministry of the Environment involve visual surveys and glue traps to capture individuals (Toda et al., 2010). However, their elusive behaviour and isolated location still causes various challenges such as low capture and detection efficiency in low-density areas and the unintended capture of non-target species, including native lizards (Hiroyama and Iwai, 2022; Obata and Iwai, 2023). Therefore, non-invasiveness should be considered in the detection method of green anole, as well as simplicity and efficacy.

Environmental DNA (eDNA) analysis is recognised as a highly sensitive method for detecting macroorganisms and is particularly effective for identifying rare and invasive species (Bell et al., 2024; Tsuji et al., 2024). However, the majority of eDNA studies to date have focused on aquatic macroorganisms, and applications for detecting terrestrial organisms remain in the early stages (Aucone et al., 2023; Gregorič et al., 2022; Lyman et al., 2022; Lynggaard et al., 2024). Newton et al. (2025) and our literature search on reptile studies identified only 76 publications up to 2024, the majority of which used water samples for eDNA detection in turtles and snakes (Fig. 1). The two previous studies that detected lizard species attempted eDNA detection from water, soil or the surface of artefacts, as target species are semi-aquatic or ground-wandering (Kyle et al., 2022; Reinhardt et al., 2019). Additionally, although not included in the above count, a most recent study published in January 2025 reported the successful detection of iguana eDNA using swab and glue tape, suggesting the potential of these methods (van Kuijk et al., 2025). However, swab and tape collection methods collect eDNA from a very limited area, which can easily lead to false negative results when targeting low population density species.

**Figure 1.**
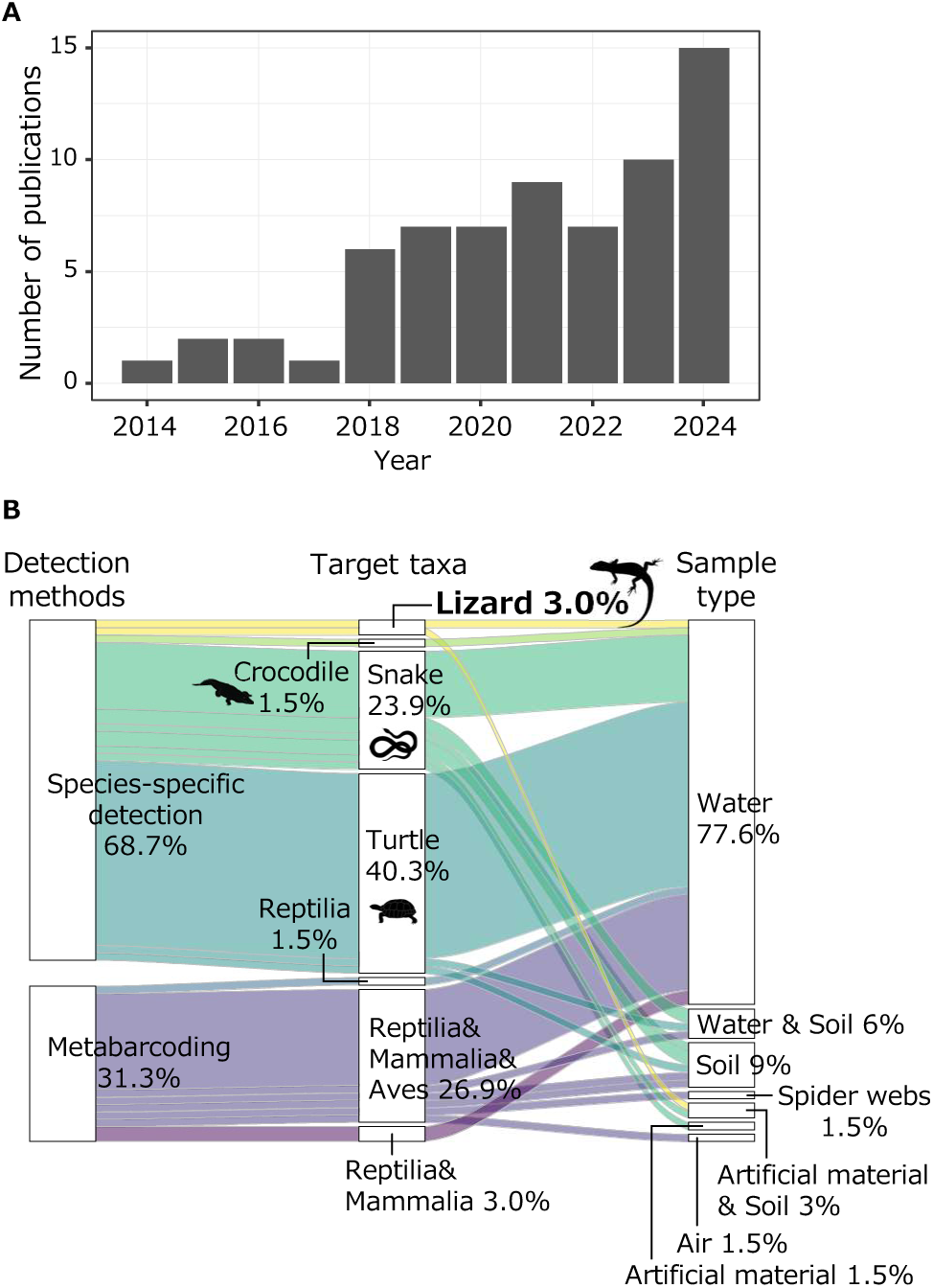
Literature search results: (A) reptiles eDNA publications per year from 2014 to 2024 (n = 67) and (B) breakdown of detection methods, target taxa and sample types used in each study

Various eDNA sampling techniques have been considered for detecting arboreal species, but there is still room for development in terms of practicality. Previous studies on arboreal species have tried various methods, including surface swabbing with rollers (Allen et al., 2023; Kyle et al., 2022; Newton et al., 2022; Valentin et al., 2020), glue tape sampling (Aucone et al., 2023; van Kuijk et al., 2025), passive rainwater sampling (Macher et al., 2023; Sakata et al., 2023) and washing off with water spray (Lee-Rodriguez et al., 2024; Valentin et al., 2020; Yoneya et al., 2023). Notably, a method using a moistened paint roller to collect eDNA from large surface areas has suggested the potential to effectively detect arthropods, mammals, and reptiles by suspending the collected eDNA in water, followed by filtration and concentration. On the other hand, the heavy and bulky equipment and supplies required for sampling, such as water, peristaltic pumps, and rollers, represent a limitation in terms of preparation effort and cost (D. L. Peterson et al., 2022; Valentin et al., 2020), which decreases the availability of methods in inaccessible area. Rainwater sampling has the advantage of collecting eDNA from a wide surface area; however, its feasibility and conditions are inherently dependent on rainfall events. In contrast, water spray aggregation actively washes and collects eDNA, providing the advantage of more easily controlled sampling conditions. Although these methods hold promise for detecting arboreal reptiles, further research is necessary to develop robust sampling methods that allow detection in low population density areas by comprehensive sampling with limited resources.

The objective of this study was to develop new survey methods for collecting and detecting terrestrial eDNA accumulated on the surfaces of tree leaves to enable the effective detection of green anole invasions. Specifically, we aimed to detect their invasions at the early stages, when densities are low, and to improve the success rate of exclusion and population expansion control. We examined the following three points: (1) comparison of eDNA detection sensitivity between wiping and spray aggregation methods, (2) relationship between eDNA concentrations and population density and (3) effectiveness of pre-filtration of sample water to improve detection sensitivity. Based on the results, we propose an efficient and cost-effective terrestrial eDNA detection method for arboreal reptiles, including the green anoles. Furthermore, we also discuss the potential in quantitative assessment and the scope for further methodological improvements. The user-controlled terrestrial eDNA surveys of forest landscapes will provide a valuable tool for increasing the success rate of monitoring and controlling biological invasions by reptiles and other terrestrial macroorganisms.

## 2 Materials and methods

### 2.1 Literature search and data compilation on Reptile eDNA detection

Following Newton et al. (2025), which organised the literature on the detection of terrestrial amniotes, that is, all tetrapods excluding the amphibians, a literature search was conducted to identify studies published in 2024 that included reptile eDNA detection. Briefly, we searched the Web of Science in March 2025 for peer-reviewed scientific articles using the following terms: “Metabarcod*” OR “Environmental DNA” OR “eDNA” AND “reptile” OR “reptilia” OR “snake” OR “lizard” OR “squamata” OR “tortoise” OR “turtle” OR “testudines” OR “crocodile” OR “alligator” OR “crocodilia” OR “tuatara” OR “rhynchocephalia” OR “terrapin” OR “skink” OR “caiman”. After manually excluding studies that did not focus on eDNA, reviews, government documents and entirely industrial documents, 15 papers remained. Additionally, from Table S2 in Newton et al. (2025), we extracted 52 papers containing reptile detection published between 2012 and 2023. For a total of 67 papers, the following information was recorded and organised: eDNA detection methods, target taxonomic groups and substrate type collected (Table S1).

### 2.2 Study site

The Ogasawara Islands consist of over 30 islands in the Pacific Ocean, approximately 1,000 km from mainland Japan (Fig. 2A). Since their formation, they have never been connected to the continent and have established a unique ecosystem with many endemic species. For this reason, the Ogasawara Islands were inscribed as a World Natural Heritage Site in June 2011 (Yoshida, 2018). The invasion of the green anole was first found in the northern part of Chichi-jima in the late 1960s (Matsumoto et al., 1980), followed by the reports in Haha-jima and Ani-jima in the early 1980s and 2013, respectively (Shimizu, 2013; Suzuki and Nagoshi, 1999). In Ani-jima, the distribution area of the green anole has continued to expand, and it is now found in almost the entire area except for the northernmost part (Yamamoto, 2024).

**Figure 2.**
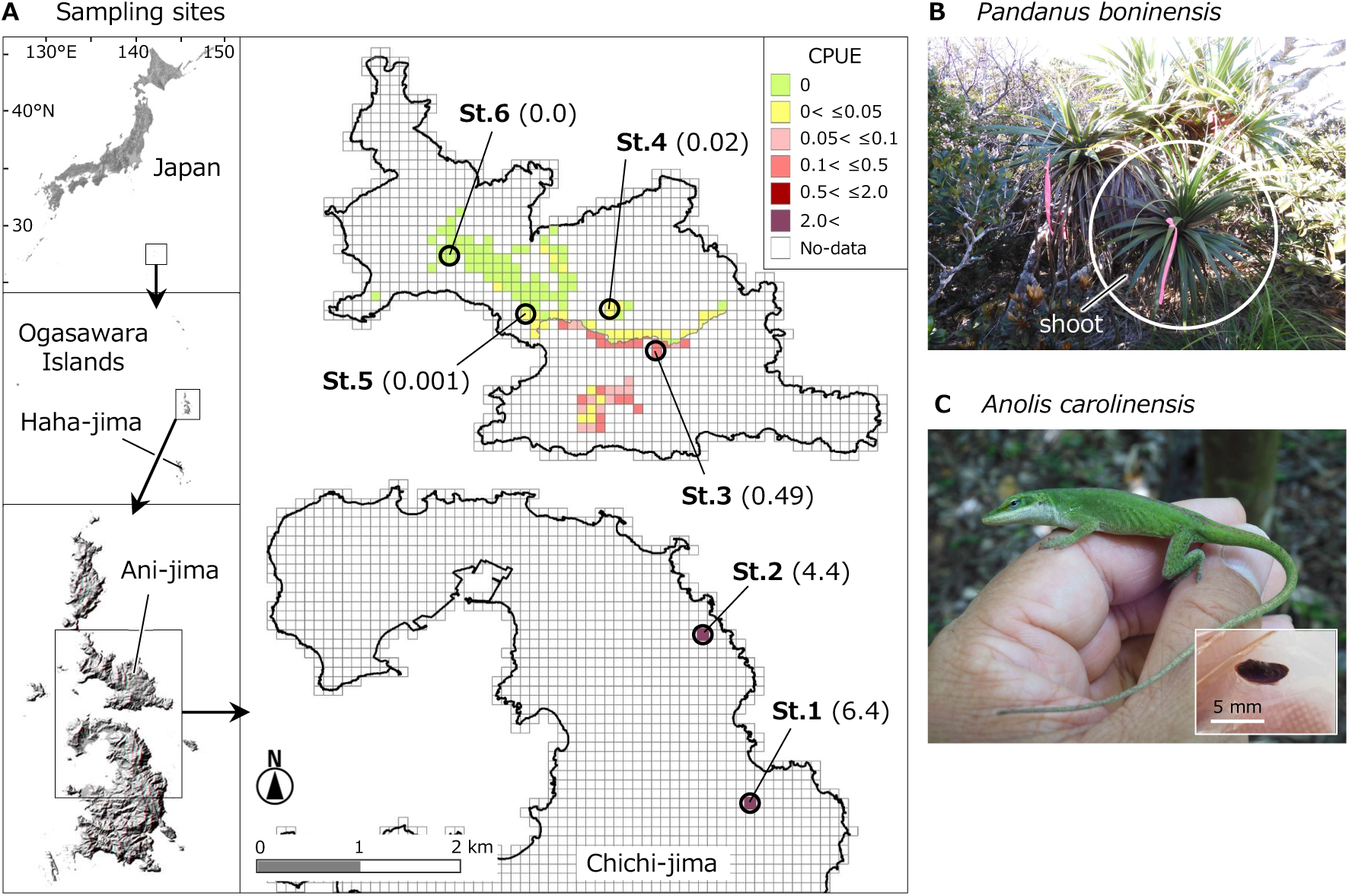
(A) Map of survey sites in the Ogasawara Islands, Japan. The numbers in brackets indicate CPUE (number of captured individuals/100 sticky traps-days) in the sticky trap-based eradication project by the Ministry of Environment. As no sticky traps were set at site 2, the CPUE estimated from the number of individuals observed per 10 minutes of line census was shown. At site 6, there were no capture records, although there were sightings of anoles, (B) *Pandanus boninensis*, (C) Green anoles and their faeces

### 2.3 Species-specific primer probe set development for green anole

The mitochondrial 16S ribosomal RNA (rRNA) sequences of the green anole and all reptiles except sea turtles inhabiting the Ogasawara Archipelago (*Lepidodactylus lugubris*, *Hemidactylus frenatus*, *Cryptoblepharus nigropunctatus* and *Indotyphlops braminus*) were downloaded from the National Center for Biotechnology Information database (NCBI; https://www.ncbi.nlm.nih.gov/) (Table S2). The obtained sequences were aligned using MAFFT (https://mafft.cbrc.jp/alignment/server/), and primers were manually designed at positions containing 3 to 5 species-specific nucleotide variations for green anole within the five bases from the 3′ end: AnolisCa_16S-F (5’-ACC CTG TGG AGC TTT AAA TTT CAA C-3’), AnolisCa_16S-R (5’-TCA ATA TTA GCT TTG GCT TGT AGG C-3’). A *in silico* test was conducted using Primer-BLAST (https://www.ncbi.nlm.nih.gov/tools/primer-blast/) with default settings to evaluate the species specificity of the designed primer set. The TaqMan probe was manually designed within the primer amplification region at a position where more than 50% of bases were mismatched with all non-target species: AnolisCa_16S-Pr (5’-[FAM]-ACA TCC GAG CAA ATA AGG CAC TGC CT-[BHQ]-3’). Additionally, an *in vitro* test was performed to validate the species specificity of the designed primer probe set using genomic DNA extracted from the green anole (five individuals) and four non-target species (three individuals per species) (Table S3). As a DNA template, 100 pg of genomic DNA was used in each PCR reaction. The real-time PCR conditions were identical to those used to analyse eDNA samples, as described below. In addition to evaluating amplification curves via real-time PCR, the presence or absence of DNA bands in the PCR products was confirmed by electrophoresis.

### 2.4 Experiment 1: Comparison of wiping and spray aggregation methods for green anole detection

In Experiment 1, the effectiveness of the (1) wiping and (2) spray aggregation methods for detecting green anole eDNA was compared. Sampling was conducted at two sites (st.1 and st.2; Fig.2A) on Chichi-jima, where they inhabit with high densities. Green anoles are frequently observed on the leaves of *Pandanus boninensis*, an endemic small evergreen tree widely distributed across the Ogasawara Islands (Aota et al., 2021; Mitani et al., 2020). Therefore, as their leaves were expected to retain traces of green anoles, such as skin fragments and faecal matter, *P. boninensis* was selected as a sampling target in this study (Fig. 2C).

*Pandanus boninensis* has about 20 to 30 large, sword-shaped leaves growing in clusters from the tip of its thick trunk, and this leafy area is called a shoot (Fig. 2B). At each site, 20 and seven trees of *P. boninensis* with at least four shoots were selected for st. 1 and st. 2, respectively. To standardise sampling conditions between the two methods as much as possible, samples for comparison were collected from four shoots, equally selected from a shared pair or combination of trees (Table S4). Sampling using each method was conducted alternately. To prevent contamination, all equipment was decontaminated before each use with 10% commercial bleach, followed by ultrapure water or was replaced with disposable alternatives. The researcher wore disposable vinyl gloves during sampling and changed them among samples. On the final day of sampling at each site, 300 mL of distilled water was filtered using the same equipment and methods as the environmental samples and treated as a field blank (FNC).

#### (1) Wiping method

In the wiping method, sterilised gauze moistened with distilled water was used to wipe the surface of *P. boninensis* leaves, suspending any eDNA considered to be attached to the leaf in the distilled water (Fig. 3A). For each sample, a single piece of gauze (22 cm × 20 cm) was used to wipe all leaves of four shoots selected from two to four *P. boninensis* trees. The gauze used for wiping was washed with 1 L of distilled water in a bucket covered with a plastic bag after each wipe of one shoot to suspend the adhering material in the water, then tightly wrung out and used for wiping the next shoot. After wiping all the leaves of one shoot, the gauze was rinsed with 1 L of distilled water in a plastic bag-covered bucket to suspend adhered substances in the water. The gauze was then wrung out and reused for wiping and resuspending the next shoot. The resuspended water was filtered using a Sterivex filter cartridge (nominal pore size = 0.45 µm, Merck Millipore, USA) and a 50 mL syringe (SS-50LZ, Terumo, Japan). The filtration volume that could be filtered by maximum effort was recorded for each sample, as the Sterivex was clogged during filtration (Table S4). To prevent eDNA degradation, 1 mL of Buffer ATL from the DNeasy Blood and Tissue kit (Qiagen, Hilden, Germany) was added to each Sterivex post-filtration (Wu and Minamoto, 2023). All Sterivex samples were then transported to a research facility in the Ogasawara Islands in a cooler box with ice packs and stored in a refrigerator at about −20 °C. After the survey schedule was completed, all Sterivex samples were transported under refrigeration to a laboratory on the mainland. The eDNA extraction and quantitative PCR were performed following the methods described below.

**Figure 3.**
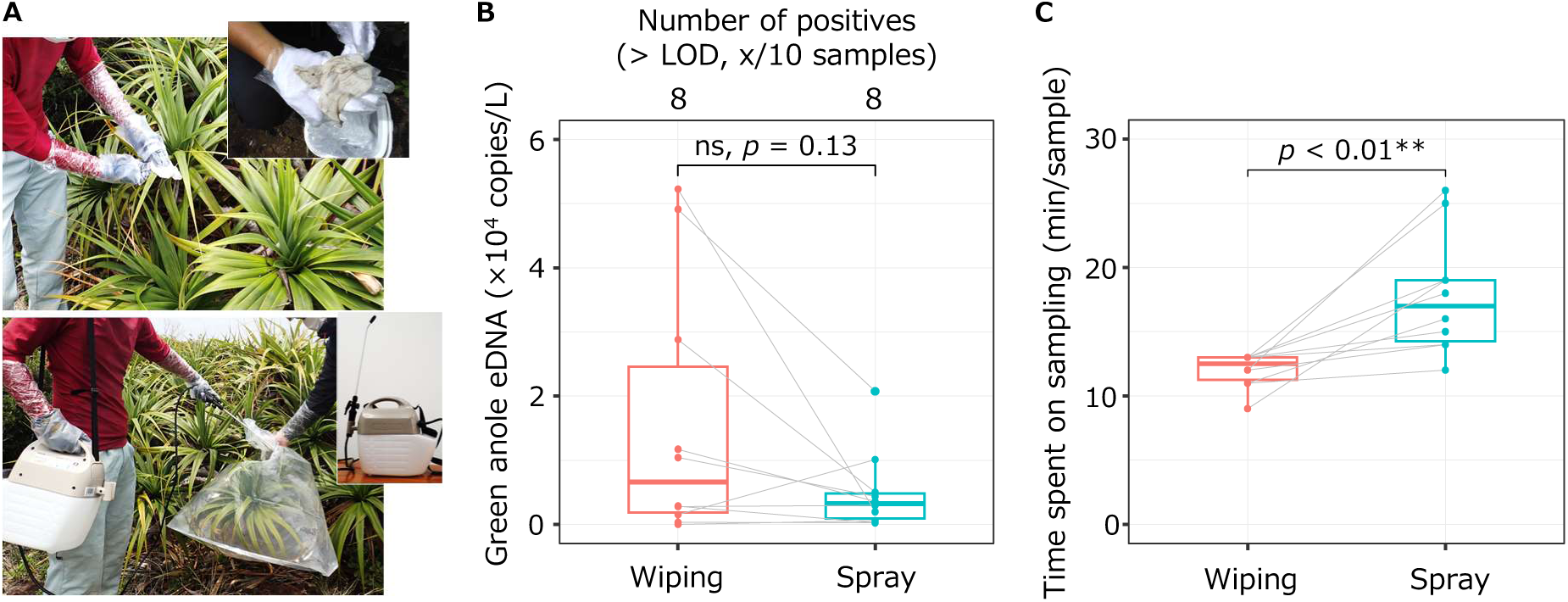
(A) eDNA sampling by wiping method (top row) and spray aggregation method (bottom row), (B) Comparison of the number of PCR-positive reactions (> LOD, x/10 samples) and detected green anole eDNA concentrations between the two methods (Wilcoxon signed-rank test, p = 0.13), (C) Comparison of the time spent on eDNA sampling in each method (from the start of sampling to the end of filtration; Wilcoxon signed-rank test, p < 0.01)

#### (2) Spray aggregation method

In the spray aggregation method, distilled water was sprayed onto the leaves of a *P. boninensis* shoot, which were covered with a plastic bag, and eDNA was collected by catching the runoff water into the bag (Fig. 3A). The spray aggregation was conducted using a battery-operated sprayer (GT-5HD-AAA, KOSHIN LTD., Japan). For each sample, 1–1.4 L of distilled water was sprayed onto a total of four shoots from 2 to 4 *P. boninensis* trees. The spray rate was 360 mL/min, and the water recovery rate after spray aggregation ranged from 41% to 92% (Table S4). The water collected in the plastic bag was filtered using the same method as the wiping method, after which 1 mL of Buffer ATL was added. The Sterivex samples were transported and stored under refrigeration at about −20 °C. As the collected sample water contained suspended particles comparable to those from the wiping method and due to the filter clogged, the filtration volume that could be filtered with maximum effort was recorded for each sample (Table S4). The eDNA extraction and quantitative PCR were performed following the methods described below.

### 2.5 Experiment 2: Relationship between the eDNA concentration detected using the wiping method and green anole density

In Experiment 2, to investigate the relationship between green anole density and eDNA concentrations, sampling by the wiping method was conducted at five sites (st. 2–6) on Chichi-jima and Ani-jima, which have different population densities of the anole. In addition, the effectiveness of pre-filtration of sample water to improve survey efficiency and detection sensitivity was also investigated. As an indicator of population density, the catch per unit effort (CPUE; number of captured individuals/100 sticky traps-days) from the Ministry of the Environment of Japan’s project to eliminate green anoles using sticky traps was used (Fig. 1A). The wiping method was adopted as demonstrated in Exp. 1 to be effective in terms of detection sensitivity and procedural simplicity (see Results).

Four *P. boninensis* trees with at least four shoots were selected at each site. Using the same procedure as in Exp.1, wiping sampling was conducted on four shoots per tree, and 1 L of distilled water in which leaf surface deposits were suspended was treated as the individual sample (Table S5). However, at st. 3, due to the insufficient number of suitable trees, one sample was collected from two shoots derived from two different trees. Each 1 L of sample water was filtered using a Sterivex filter unit to the volume that could be filtered with maximum effort in the same way as in Exp. 1 and the filtration volume was recorded. The remaining sample water was pre-filtered using a commercial coffee filter (KANAE Co., Ltd., Ehime, Japan), followed by filtration with a Sterivex filter unit to the volume that could be filtered with maximum effort in the same way as in Exp. 1 and the filtration volume was recorded. Following sample collection at st. 2, st. 3 and st. 5, 300 mL of pre-filtered distilled water was filtered using the same equipment and methods as the environmental samples and treated as an FNC. After filtration, 1 mL of Buffer AL was immediately added to the Sterivex. All Sterivex samples were transported and stored under refrigerated conditions following the same flow as in Exp.1. During field sampling, the same contamination control measures as in Exp. 1 were implemented and maximum care was taken to minimise the risk of contamination. For data analysis, the quantified eDNA copy number of green anole was converted to a concentration per litre of water used for suspending.

### 2.6 DNA extraction from Sterivex filter unit samples

Environmental DNA was extracted from each Sterivex filter unit using a DNeasy Blood & Tissue Kit according to the method described by Wu and Minamoto (2023) with some modifications. Since 1 mL of Buffer ATL had been added to each Sterivex filter unit immediately after filtration, an additional 50 µL of Proteinase K was added. The filter units were then incubated at 56 °C for 30 min while being rotated using a rotator (RotoFlex R2200, Cole-Parmer, Vernon Hills, USA). After incubation, the inlet cap (VRMP6) was removed and replaced with a 5-mL tube (cat no. 67-2395-22, AS ONE, Osaka, Japan). The Sterivex was centrifuged with the inlet side down at 6,000 g for 1 min. The filtrate collected in the 5-mL tube was mixed with 1 mL of Buffer AL by pipetting and re-incubated at 56 °C for 10 min. Subsequently, 1 mL of 99.9% ethanol was added to the mixture, and the entire was passed through the DNeasy Mini Spin Column in multiple steps. The subsequent DNA purification steps followed the manufacturer’s instructions. Finally, DNA was eluted from the column with 100 µL of Buffer AE. The eDNA extraction process was conducted separately from the real-time PCR experiments.

### 2.7 Quantitative real-time PCR Assay

Quantitative real-time PCR (qPCR) was performed in triplicate per sample using the Light Cycler 96 system (Roche, Basel, Switzerland). The composition of the qPCR reaction solution (total volume, 15 µL) was 900 nM of each primer (AnolisCa_16S-F/R), 125 nM of the TaqMan probe (AnolisCa_16S-Pr), and 2.0 µL of DNA template in 1× TaqMan Environmental Master Mix (Thermo Fisher Scientific). In all qPCR runs, a dilution series of known concentrations (3×10^1^, 3×10^2^, 3×10^3^ and 3×10^4^ copies per reaction) and PCR negative controls (PCR-NCs, ultrapure water) were also amplified alongside environmental samples. The qPCR thermal cycling conditions were as follows: 50 °C for 2 min, 95 °C for 10 min, followed by 55 cycles of 95 °C for 15 s and 60 °C for 60 s. Although detected at low concentrations (below the LOD), 0.4 and 0.5 copies/qPCR reaction were detected in the PCR-NC in PCR-Run 01 for Exp.1 and FNC in st.2 for Exp.2. To address the effect of contamination occurrence, the DNA copy numbers detected from each negative control were subtracted from the corresponding samples (sample IDs 1 to 14 in Exp.1 and all samples of st. 2 in Exp.2), and subsequent data analysis was performed.

Furthermore, we estimated the limit of quantification (LOQ) and limit of detection (LOD) using 15, 10, 6, 5, 4, 3, 2 and 1 DNA copies as templates per reaction. The LOD was set to the lowest concentration at which positive reactions were obtained in two or more of the three PCR replicates. For the LOQ, a concentration was adopted that fulfilled the following two conditions: 1, positive reactions were obtained in all three PCR replicates; 2, the R² value was greater than 99.0 when a standard line was drawn together with four dilution series. The R² values for the standard curves of all qPCR assays in this study ranged from 0.99 to 1.00. For data analysis, estimated eDNA copy numbers were converted to DNA concentration per litre of water suspended or sprayed.

### 2.8 Inhibition test

The occurrence of PCR inhibition in each sample was assessed using DNA adjusted to known concentrations of *Plecoglossus altivelis*, which is absent from the Ogasawara Islands. The species-specific primer probe set for *P. altivelis* was developed by Yamanaka and Minamoto (2016), and the sequence is as follows: Paa-cytb-F (5’-CCT AGT CTC CCT GGC TTT ATT CTC T-3’), Paa-cytb-R (5’-GTA GAA TGG CGT AGG CGA AAA-3’), Paa-cytb-Pr (5’-[FAM]-ACT TCA CGG CAG CCA ACC CCC-[BHQ]-3’). Reagent preparation for qPCR was conducted as described above, with an additional 20 copies of *P. altivelis* DNA per reaction added to the reaction mix. All other qPCR condition settings are the same as for green anoles. PCR inhibition was considered to have occurred if the quantified copy number was 50% below 20 copies (i.e., when the Cq value was delayed by 1 cycle). Samples in which PCR inhibition was recognised were corrected by dividing the quantified copy number of green anole by the ratio of detected *P. altivelis* copy number (x/20 copies), and the corrected values were used for data analysis.

### 2.9 Statistical analysis

All statistical analyses and graphic illustrations for this study were conducted using R version 4.2.2 software (R Core Team, 2023). The α level of significance was set at 0.05. In Exp. 1, we performed a Wilcoxon signed-rank test to compare the eDNA concentration between the wiping and spray aggregation methods by the exactRankTests package ver. 0.8.34. Similarly, the time taken from the start of the eDNA collection to the end of filtration by each method was compared between the two methods using the Wilcoxon signed-rank test. In Exp. 2, generalised linear model (GLM) analysis was performed to investigate the relationship between eDNA concentration and CPUE (number of captured individuals/100 sticky traps-days) in green anoles. The probability distribution of the response variable (eDNA concentration) followed a Poisson distribution with the log-link function. Furthermore, to investigate the mitigating effect of pre-filtration on PCR inhibition, the detected concentrations of artificially added *P. altivelis* DNA were compared using a paired t-test between with and without pre-filtration.

## 3. Results

Since the first publication on reptile eDNA detection in 2014, 76 relevant papers have been published to date, with the number of publications showing a gradual annual increase (Fig. 1A). In 2024, 15 papers were published, marking the highest number in the past decade. Detection methods were categorised into species-specific detection and metabarcoding (Fig. 1B). In species-specific detection, the majority of studies focused on turtles and snakes, with only two studies targeting lizards (Table S1; Kyle et al., 2022; Reinhardt et al., 2019). Furthermore, water samples were used for eDNA collection in 83.6% of the studies.

An *in silico* test using Primer-BLAST showed that the designed primer set could specifically detect the green anole at the study sites in the Ogasawara Islands. The amplicon length was 151 bp, with melting temperatures (Tm) of 60.2 °C and 59.7 °C for the forward and reverse primers, respectively. The Tm of the TaqMan probe was 66.8 °C. In an *in vitro* test using genomic DNA, species-specific and efficient DNA amplification were confirmed by qPCR and electrophoresis of the PCR products. The LOD was set at 2 copies per qPCR reaction, with a positive reaction in 2/3 qPCR replicates (Table S6). The LOQ was set to 10 copies per qPCR reaction (Table S6).

Green anole eDNA was detected using both wiping and spray aggregation methods. No samples showed PCR inhibition. Of the 10 samples from each method, eight samples were detected above the LOD in both methods (Table S7, Fig. 3B). Excluding three sample pairs where concentrations were below the LOD for both methods, eDNA concentrations were compared between methods for the remaining seven sample pairs, and no significant differences were found (Wilcoxon signed-rank test, p = 0.13; Fig. 3B). The time required for eDNA sampling was 12.0 ± 1.3 and 17.8 ± 4.7 (mean ± SD) minutes for the wiping and spray aggregation methods, respectively, with the wiping method taking less time (Wilcoxon signed-rank test, p < 0.01; Fig. 3C).

A significant positive relationship was found between eDNA concentration and green anole CPUE (GLM, p < 0.01; Fig. 4). This relationship was also observed when samples with eDNA detected above the LOQ were only used in the analysis (GLM, p < 0.01). Samples where the detected concentration did not exceed the LOQ or LOD tended to be more prevalent at sites with lower CPUE.

**Figure 4.**
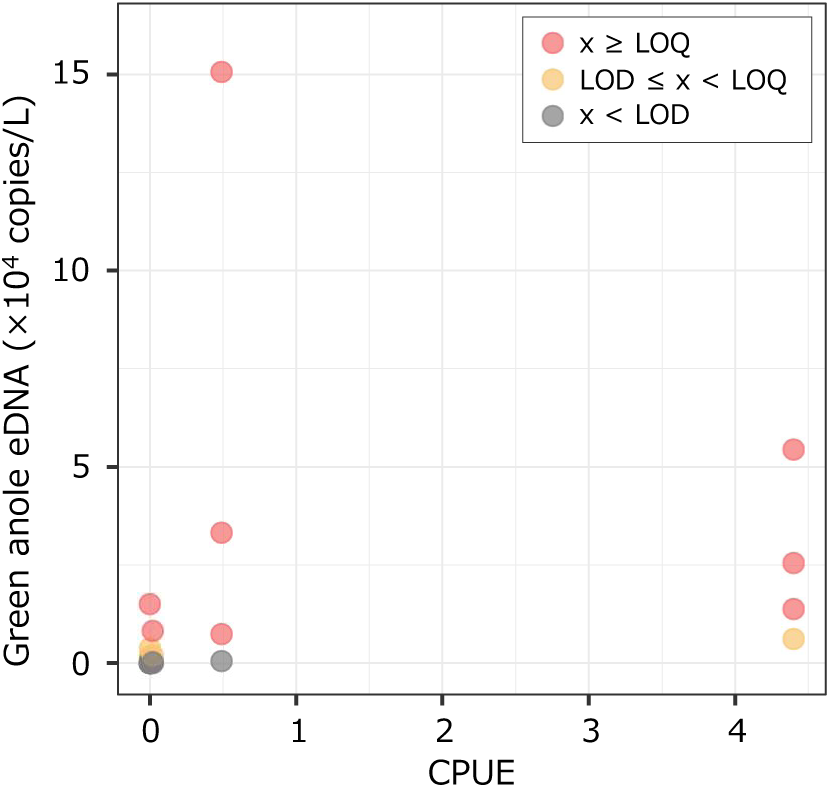
Relationship between CPUE (number of captured individuals/100 sticky traps-days; as a surrogate for habitat density) and detected eDNA concentrations of green anoles at each study site (GLM, p < 0.01)

To address filter clogging, the effect of pre-filtration with coffee filters on green anole eDNA detection and PCR inhibition levels was investigated. At sites with medium or higher population densities (st. 2, 3, and 4), pre-filtration decreased the number of samples with eDNA detected above the LOQ by one to two samples (Fig. 5A). In the inhibition test, *P. altivelis* DNA should be detected at 20 copies/qPCR reaction in the absence of inhibition. However, with and without pre-filtration, the detected *P. altivelis* DNA concentrations were lower than the expected 20 copies per qPCR reaction (without pre-filtration, 14.0 ± 6.4 (mean ± SD) copies/qPCR reaction; with pre-filtration, 14.4 ± 2.7 copies/qPCR reaction: Table S8). No significant differences were observed in the detected *P. altivelis* DNA concentrations between with and without pre-filtration (paired t-test, p = 0.74; Fig. 5B). However, in some samples, the detected *P. altivelis* DNA copies were less than 50%, so PCR inhibition was considered to have occurred, and the quantitative values for green anole were corrected (Table S8).

**Figure 5.**
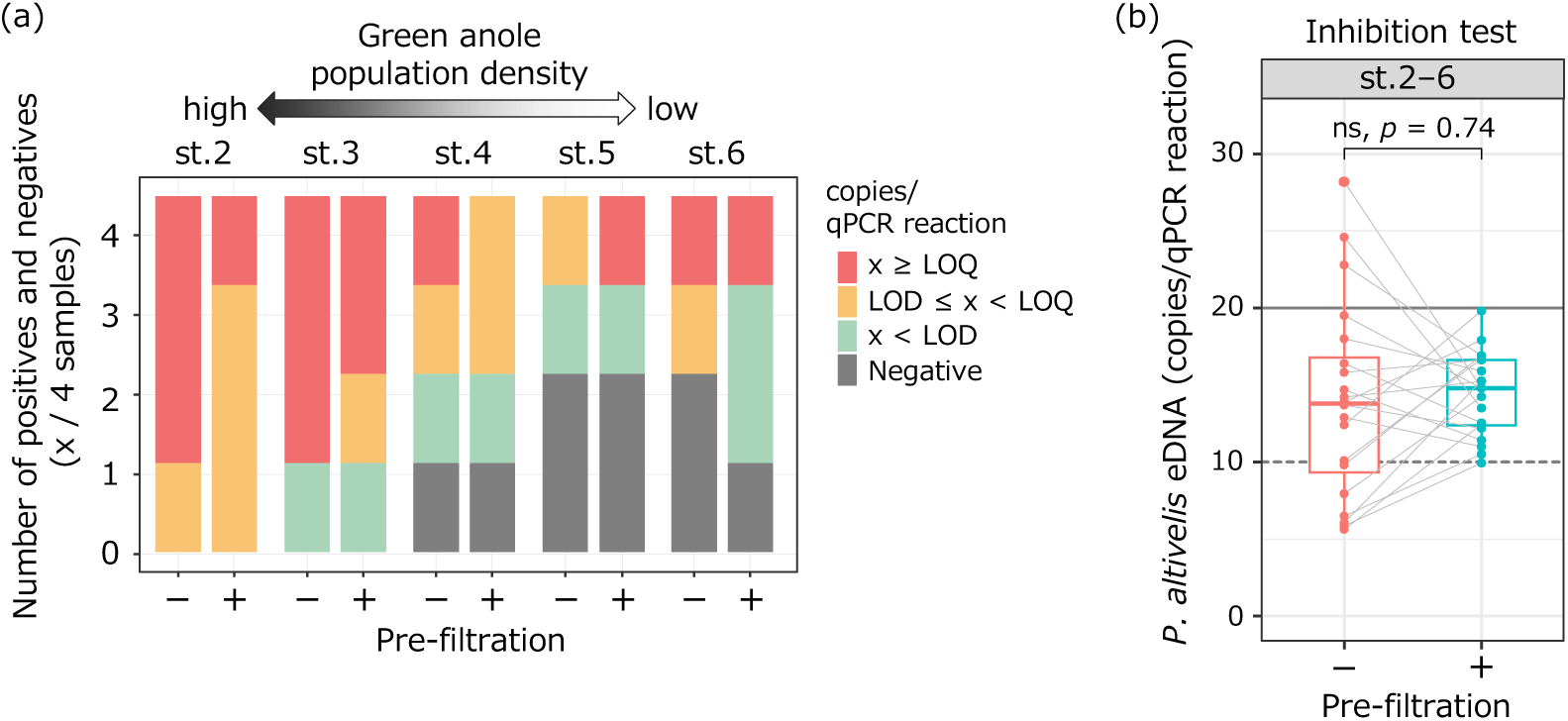
(A) Comparison of the number of PCR positive and negative reactions between with and without pre-filtration for each site, (B) Results of the inhibition test using *P. altivelis* DNA between with and without pre-filtration (paired t-test, p = 0.74). The solid and dashed lines indicate the 20 copies added to each qPCR reaction and the 10 copies set as the criterion for determining that PCR inhibition had occurred, respectively

## 4. Discussion

In this study, we present the first evidence that terrestrial eDNA collected from leaf surfaces can be used effectively to detect the arboreal invasive lizard, the green anole. Notably, positive detections were obtained even from sites with extremely low population densities (sites 5 and 6). As suggested by numerous previous studies using aquatic eDNA (Rourke et al., 2022), when the collection range is clearly defined, terrestrial eDNA concentrations could also be used as an indicator of the habitat density of the target species. Our findings suggest that the application and further development of eDNA analysis offer a promising new approach for monitoring invasions of often-overlooked reptiles. This study provides several key insights into the operationalisation of eDNA techniques for surveying arboreal invasive reptiles.

The detection of species with low population densities is inherently challenging. In such cases, eDNA-based surveys, which combine high efficiency and detection sensitivity to achieve comprehensive sampling with minimal effort, are particularly suitable. In Experiment 1, we attempted to detect eDNA from leaves using two methods: the wiping method and the spray aggregation method, both of which successfully detected the green anole without relying on visual detection. Furthermore, both methods showed positive results in 80% (8/10 samples), indicating comparable detection sensitivity between the two methods. Previous studies by Kissling et al. (2018) and van Kuijk et al. (2025), which employed swabbing methods using small swabs, collected samples from visually identified positive surfaces such as objects or perches in the breeding room, but reported lower positivity rates (57.0% and 10.5%, respectively). The success of terrestrial eDNA-based surveys is determined by collecting samples from positions where the target species shed DNA and by aggregating as much eDNA as possible (Kyle et al., 2022). Therefore, the ability to effectively sample the entire leaf surface, which is frequently utilised by green anoles, likely explains the high detection sensitivity observed in the wiping and spray aggregation methods. On the other hand, in terms of sampling simplicity, the swabbing method was found to be superior to the spraying method, as it allowed surveys to be conducted in less time and with fewer materials. The simplicity of sampling is crucial for the feasibility of multi-site and high-frequency monitoring surveys, concerning the future application of this method in Ogasawara. Furthermore, the use of disposable gauze for surface wiping has a significant advantage by keeping consumable costs and contamination risks relatively low. Consequently, we recommend the wiping method as an effective method for detecting arboreal reptiles, including the green anole.

The significant positive relationship observed between eDNA concentrations of the green anole and its population density in Experiment 2 raises expectations for quantitative terrestrial biodiversity assessment in future terrestrial eDNA studies. Since the detection of macroorganisms using eDNA began, the quantitative assessment of target species based on eDNA concentrations has remained a prominent topic of discussion for over a decade (Blackman et al., 2024). In studies targeting fish, 90% of studies have shown a positive correlation with biomass (Rourke et al., 2022). However, as terrestrial eDNA research is still relatively new, few studies have investigated the relationship between eDNA concentration and quantitative information, such as species population density. Therefore, continued research to accumulate the information on this relationship is crucial; simultaneously, efforts to reduce observational noise would be necessary. One of these is to understand and unify the spatial range from which eDNA is collected within each study. When collecting eDNA from air or rain, it is expected that the complexity of meteorological factors, such as wind (Newton et al., 2024), may influence the amount or the collection range of eDNA, which is uncontrollable and difficult to consider. In contrast, surveys using the wiping method allow the eDNA collection area to be freely set and standardised among samples, which may have led to the positive results in this study. This methodological advantage is also shared by spray aggregation, roller, and swab methods. Standardising and understanding the eDNA collection range would strengthen the link between the density and/or biomass of the target species and the probability of recovering eDNA, and would facilitate quantitative assessments based on eDNA concentrations.

Environmental factors at sampling sites are suggested to play a crucial role in the persistence and detection success of eDNA (Barnes and Turner, 2016), and particular attention should be paid to temperature, ultraviolet (UV) exposure, and rainfall in terrestrial environments (Newton et al., 2025; Valentin et al., 2021). Terrestrial eDNA on leaf surfaces is exposed to stronger UV radiation and higher temperatures than eDNA in aquatic environments, meaning eDNA degradation rate can vary significantly depending on sunlight conditions. Furthermore, rain and fog have been shown to temporarily wash away eDNA from leaf surfaces, as supported by reports of eDNA detection from rainwater (Macher et al., 2023; Sakata et al., 2023; Valentin et al., 2020). In Experiment 2 of this study, eDNA sampling was carried out by selecting *P. boninensis* under the same sunlight conditions as far as possible. However, as weather conditions cannot be controlled, there was a small amount of rainfall a few minutes before sampling for st. 5 and the day before sampling for st. 2. Although there is no definitive evidence, it is possible that eDNA detection below the LOQ (1/4 sample) was affected by rainfall, at least for the densely inhabited st. 2. Eliminating or controlling these natural factors is extremely difficult, but when scheduling surveys, avoiding periods of high temperature or intense UV radiation, and considering rainfall conditions in the days before sampling, could help enhance the success of the research.

For sampling using the wiping method, further method improvements to mitigate filter clogging by suspended solids and the inclusion of inhibitors could contribute to improvements in eDNA yield, detection sensitivity, and reproducibility. The pre-filtration method, commonly used to selectively remove large suspended matter other than eDNA from water samples, was employed in this study (Robson et al., 2016; Stoeckle et al., 2020). Coffee filters, which have a proven track record in fish eDNA, were used as pre-filters (Takasaki et al., 2021), but contrary to our expectations, they did not improve the sensitivity of green anole detection or reduce PCR inhibition. This may have been due to some green anole eDNA being captured along with suspended matter in the coffee filter. There is no information on the particle size of terrestrial eDNA, but eDNA released from reptiles is probably shed mainly as faeces and skin fragments, and it is therefore expected to be larger than that of fish (eDNA particle size in fish, 0.7 μm to 8 μm; Tsuji et al., 2022; Tsuji and Shibata, 2025). It is also possible that the main suspended particles causing the filter clogging were not soil dust, which is rich in PCR inhibitors, but rather fine fibrous hairs or colonies of sooty mould on the leaf surface. This is supported by the fact that no change in the PCR inhibition level was observed after pre-filtration and by the presence of fibrous materials and black particles on the filter after filtration. In particular, when targeting species with low population densities, increasing the filtration volume while mitigating PCR inhibition improves detection probability and quantifiability, so it is worth continuing to improve the method for this aspect in the future.

This study examines two methods for detecting the arboreal invasive lizard, the green anole, in terms of detection sensitivity and simplicity, highlighting the potential utility of wiping methods. Environmental DNA analysis has generally been considered unsuitable for detecting non-aquatic reptiles, and there has been limited research in this area. However, the increased use of terrestrial eDNA will rapidly accelerate its application to the detection of reptiles shortly. Furthermore, as arboreal environments are major habitats not only for reptiles but also for other taxonomic groups, the benefits of using eDNA from leaf surfaces are expected to extend widely to other terrestrial taxa. Given the increasing opportunities for invasion by non-native species and the rapid global loss of biodiversity, this study would attract significant interest from managers, conservationists, and researchers working with a wide range of terrestrial organisms.

## Data Availability Statement

Full details of the qPCR results for each experiment of the present study are available in the supporting information (Table S6, S7 and S8).

## Acknowledgements

We thank N. Tsuji, Y. Hisasue, H. Nagano and T. Matsumoto for their advice in planning and conducting the field survey. We are grateful to the Kanto Regional Environment Office, Ministry of the Environment, Japan (Tokyo, Japan) for allowing us to use the samples collected from the Ogasawara Islands by the project of the Ministry of the Environment in Japan.

## Statements & Declarations

### Authors’ contributions

Conceptualisation: All authors; Field survey: YM, MT, YY, KA, TN and NY; Molecular analysis: ST; Data analysis: ST, YM, MT and YY; Writing - original draft preparation: ST; Writing - review and editing: All authors.

### Compliance with Ethical Standards

All experiments were performed with attention to animal welfare.

